# Differential serotonin uptake mechanisms at the human maternal-fetal interface

**DOI:** 10.1101/2021.01.07.425725

**Authors:** Petra Baković, Maja Kesić, Marina Horvatiček, Meekha George, Maja Perić, Ivona Bečeheli, Lipa Čičin-Šain, Gernot Desoye, Christian Wadsack, Ute Panzenboeck, Jasminka Štefulj

## Abstract

Mechanisms regulating serotonin (5-HT) homeostasis at the maternal-fetal interface are important for proper placental functioning and fetal (neuro)development. Here we studied 5-HT uptake mechanisms in human primary trophoblasts, feto-placental endothelial cells and cord blood platelets, all isolated directly after birth. Trophoblasts and cord blood platelets demonstrated high-affinity 5-HT uptake with similar Michaelis constant (*Km*) values (0.60±0.27 and 0.65±0.18 μM, respectively). In contrast, feto-placental endothelial cells displayed saturation kinetics only over the low-affinity range of 5-HT concentrations (*Km*=782±218 μM). 5-HT uptake into trophoblasts was inhibited by various psychotropic drugs targeting high-affinity serotonin transporter (SERT), and into feto-placental endothelial cells by an inhibitor of low-affinity transporters. *SERT* mRNAs were abundant in trophoblasts, but sparse in feto-placental endothelial cells; the opposite was found for transcripts of the low-affinity plasma membrane monoamine transporter (PMAT). These results show for the first time the presence of functional 5-HT uptake systems in feto-placental endothelial cells and fetal platelets, cells in direct contact with the fetal blood plasma. Data also emphasize sensitivity of 5-HT transport into trophoblasts, cells facing maternal blood, to various psychotropic drugs. The multiple, high- and low-affinity, systems present for cellular 5-HT uptake highlight the importance of maintaining 5-HT homeostasis at the maternal-fetal interface.

## INTRODUCTION

Serotonin (5-hydroxytryptamine, 5-HT), a multifunctional biogenic monoamine, is best known as the brain modulatory neurotransmitter involved in mood, sleep and appetite regulation as well as in many mental health conditions. In addition, 5-HT acts as an endocrine, paracrine or autocrine agent regulating different peripheral functions such as vascular tone, hemostasis, intestinal motility, immune response, bone remodeling and energy metabolism^1,2^. During pregnancy, 5-HT mediates fine-tuning of embryonic/fetal development^3^ and plays important roles in placental physiology^4^. It modulates a number of cellular processes in placental cells, including proliferation, cell viability, cell cycle progression, apoptosis and estrogen production^5–9^. As a potent vasoactive autacoid, 5-HT also influences utero-placental and feto-placental blood flow^10,11^. In addition, early in pregnancy the placenta supplies the developing fetus with maternal and/or placenta-derived 5-HT required for a proper fetal (neuro)development^12–14^. This wide range of functions may account for various studies associating altered placental 5-HT homeostasis with pregnancy disorders such as preeclampsia^15^, fetal growth restriction^16^ and gestational diabetes mellitus^17^ as well as with mental health implications in the offspring^18–22^.

5-HT exerts its physiological effects by interacting with 14 distinct plasma membrane-bound 5-HT receptors coupled to diverse intracellular signaling pathways^23^. In addition, it regulates some physiological functions in a receptor-independent mode, by directly binding to different extracellular and cytosolic proteins^24^. New findings demonstrate that 5-HT attaches also to histone proteins and consequently participates in transcriptional regulation^25^. All these effects depend on the availability of 5-HT in the immediate vicinity of its molecular targets. Systems for transport of 5-HT across biological membranes efficiently control local 5-HT concentrations and thus play a central role in governing 5-HT actions.

There are two distinct plasma membrane transport systems for 5-HT, uptake-1 and uptake-2, characterized by high-affinity / low-capacity and low-affinity / high-capacity, respectively, kinetic properties. Uptake-1 system for 5-HT is represented by the serotonin transporter (SERT), a highly specific high-affinity 5-HT carrier, best known for mediating reuptake of 5-HT into presynaptic neurons^26^. Due to its central role in the clearance of 5-HT from the synaptic cleft, SERT is a main target of many psychotropic drugs, including widely prescribed tricyclic antidepressants (TCAs) and selective serotonin reuptake inhibitors (SSRIs). In adults, SERT also represents a major mechanism responsible for regulating active levels of 5-HT in the blood plasma, by mediating uptake of 5-HT into platelets^27^. An analogous 5-HT uptake mechanism in the fetal circulation, however, has not yet been described. Unlike SERT, carriers of the uptake-2 system, such as the plasma membrane monoamine transporter (PMAT) and organic cation transporter 3 (OCT3), exhibit low-affinity transport kinetics and readily transport 5-HT as well as other organic monoamines^28–31^.

The placenta is a complex, heterocellular organ interposed between maternal and fetal blood. Beside others, two principal cell types constituting the maternal-fetal blood barrier are trophoblasts covering the surface of villous trees and facing the maternal blood, and feto-placental endothelial cells forming the feto-placental vessels and being in direct contact with the fetal blood. There is evidence of placental uptake of 5-HT from both maternal and fetal circulation at the term of human pregnancy. A high-affinity uptake of 5-HT from maternal blood has been firstly suggested by 5-HT transport studies in plasma membrane vesicles isolated from human term placentas^32^. An analogous high-affinity 5-HT uptake system, sensitive to TCAs and SSRIs, was described in human trophoblast-like choriocarcinoma cell line JAR^33^, but to our knowladge, has not been shown in primary trophoblasts. Regarding placental uptake of 5-HT from the fetal blood, a low-affinity 5-HT uptake mechanism has been recently suggested by *in situ* dual perfusion system in rat term placentas and 5-HT transport studies in plasma membrane vesicles isolated from human term placentas^34^.

Despite the growing interest in the feto-placental 5-HT system and various expression studies, kinetic and pharmacological properties of 5-HT uptake in primary trophoblasts have not been studied. There is also a clear lack of functional studies on potential mechanisms of 5-HT uptake into cells that are in direct contact with the fetal plasma, such as feto-placental endothelial cells and fetal platelets. We hypothesized that these cells have functional systems for active uptake of 5-HT. This is important because such systems will contribute to regulation of fetal circulating 5-HT levels. Therefore, to expand understanding of the mechanisms that regulate 5-HT homeostasis during human fetal development, we examined and characterized 5-HT uptake activity in primary trophoblasts, feto-placental endothelial cells and cord blood platelets, all isolated from the human tissues directly after birth.

## RESULTS

### 5-HT uptake into primary placental cells

Human primary trophoblasts (Figure 1A) and feto-placental endothelial cells (Figure 1B) both took up 5-HT in a time- and temperature-dependent manner, demonstrating the presence of specific, carrier-mediated process. However, specific uptake measured at a submicromolar (0.1 μM) 5-HT concentration was by up to 2.9-fold higher in trophoblasts than in feto-placental endothelial cells (Figure 1C). Further, 5-HT uptake into trophoblasts was totally abolished by a high concentration (0.1 mM) of citalopram, a potent and most selective blocker of the SERT^35^ (Figure 2), demonstrating a critical role of the high-affinity 5-HT transporter in trophoblasts. In contrast, 5-HT uptake into feto-placental endothelial cells was not affected by citalopram (Figure 2) but was sensitive to increasing concentrations of decynium 22 (Figure 3), a common inhibitor of the uptake-2 (PMAT/OCT3) transporters^28–31^. These data indicated the role of uptake-2 carrier/s and undermined the involvement of SERT in 5-HT uptake into feto-placental endothelial cells. To support pharmacological data, we performed comprehensive kinetic studies measuring the initial rates of 5-HT uptake across multiple substrate concentrations, covering a range typical for high-affinity (uptake-1) and low-affinity (uptake-2) transport systems. As expected, trophoblasts (Figure 4A) demonstrated saturation curves over the high-affinity range of 5-HT concentrations (0.1 to 3.2 μM). In contrast, feto-placental endothelial cells (Figure 4B) displayed saturation kinetics compliant with the Michaelis-Menten model only over the low-affinity range of 5-HT concentrations (94 to 3000 μM). The best-fit values of Michaelis affinity constant (*Km*) and maximal transport velocity (*Vmax),* calculated in all subjects tested, are shown in Figure 4C and Supplementary Table S1. In all subjects tested, *Km* values (mean ± SD, n=9) in trophoblasts (0.640 ± 0.266 μM) were characteristic of the high-affinity^26^ and in feto-placental endothelial cells (782 ± 218 μM) of the low-affinity^28–31^ transport systems.

**Figure 1.**
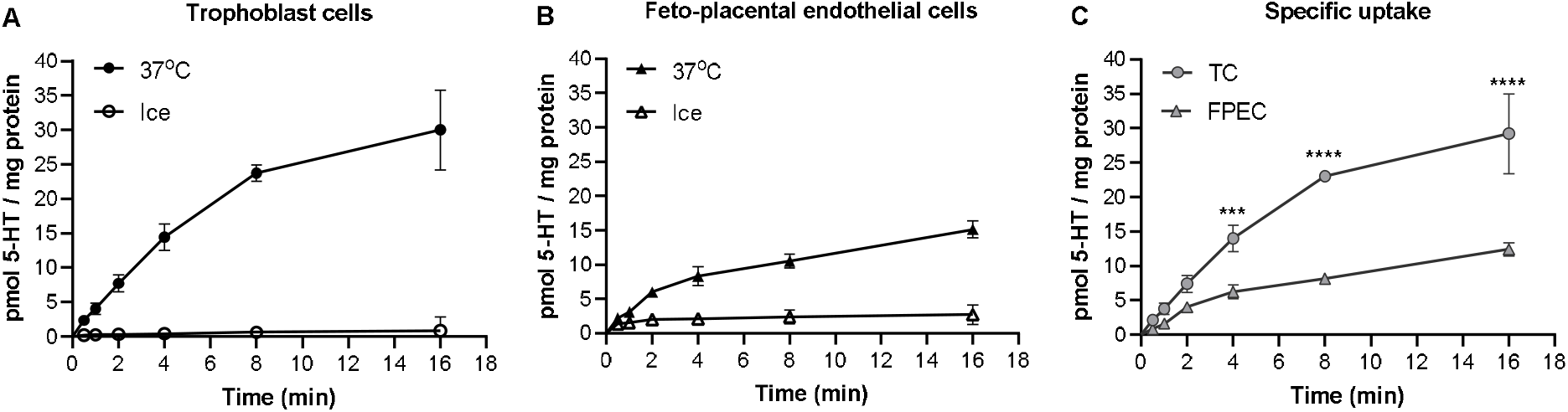
Time- and -temperature dependent uptake of 5-HT into human primary placental cells. Human primary (**A**) trophoblast cells (TC) and (**B**) feto-placental endothelial cells (FPEC) were incubated at 37°C or on ice in the presence of ^3^H-5-HT (10^−7^ M) for 0.5 to 16 min. (**C**) Specific (carrier-mediated) uptake was calculated as the difference between transport at 37°C and on ice. Shown are means ± SD (n=3). \*\*\**p*<0.0001, \*\*\*\**p*<0.0001 (Sidak’s multiple comparisons test following 2-way ANOVA).

**Figure 2.**
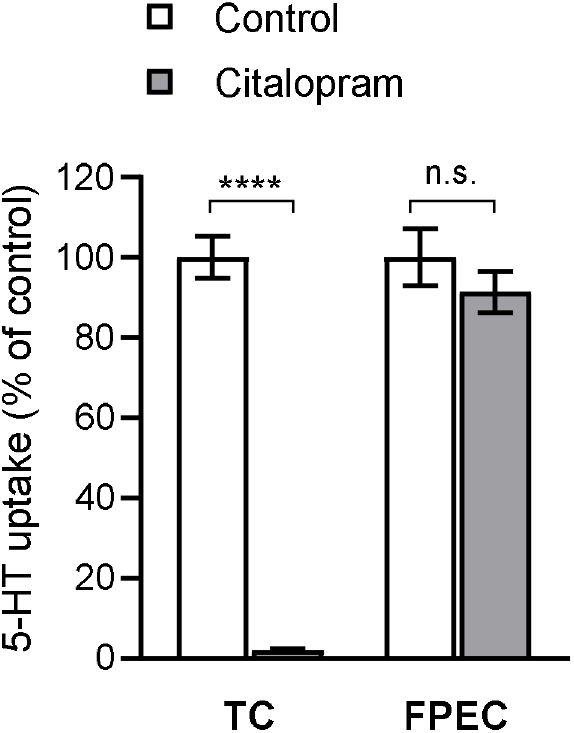
The effect of citalopram, selective inhibitor of the serotonin transporter, on 5-HT uptake into human primary placental cells. Trophoblast cells (TC) and feto-placental endothelial cells (FPEC) were incubated in the presence of ^3^H-5-HT (10^−7^ M) and citalopram (10^−4^ M) for 10 min. Values are expressed as a percentage of the control (vehicle without the drug). Data are means ± SD (n=3) and shown is representative of two separate experiments. \*\*\*\**p*<0.0001 compared to vehicle, n. s. not significant (Sidak’s multiple comparisons test following 2-way ANOVA).

### Expression of 5-HT-regulating genes in primary placental cells

To identify molecular player/s accounting for decynium-22-sensitive (Figure 3) low-affinity (Figure 4B) 5-HT uptake into feto-placental endothelial cells, we examined expression of *PMAT* and *OCT3* mRNAs in these cells, using RNA from total placental tissue as a positive control. While *PMAT* transcripts were detected in both total placental tissue and feto-placental endothelial cells, *OCT3* mRNAs were highly expressed in total placental tissue but were absent in feto-placental endothelial cells as demonstrated by absence of specific products in end-point RT-PCR analysis (Figure 5A). These data indicate that PMAT, and not OCT3, mediates uptake of 5-HT into feto-placental endothelial cells. To further support pharmacological and kinetic findings on different 5-HT uptake systems in trophoblasts and feto-placental endothelial cells, we also compared levels of *SERT* and *PMAT* mRNAs between these two cell types. Consistent with 5-HT uptake results, trophoblasts were rich in *SERT* mRNA, as demonstrated by low Cq values obtained in RT-qPCR analysis (Figure 5B). In contrast, feto-placental endothelial cells isolated from 12 different placentas yielded variable (range 26.8 to 36.7), but generally high (mean 31.2) Cq values. Relative levels of *SERT* mRNA normalized to *ACTB* mRNA (Figure 5C, left section) differed by up to 400-fold between fetoplacental endothelial cells isolated from different placentas and were on average 1000-fold lower (*p*<0.0001) in feto-placental endothelial cells than trophoblasts. This accords with earlier immunohistochemical studies reporting only a weak staining for SERT protein occasionally observed on the feto-placental endothelium of the human term placentas^14,36^. Findings of only sporadic expression of low levels of SERT, together with results of no effect of SERT blocker on 5-HT uptake (Figure 2), strengthen conclusion that this transporter is not essential in the feto-placental endothelial cells. Contrary to *SERT, PMAT* mRNA levels were about 300-fold higher (*p*<0.0001) in feto-placental endothelial cells than in trophoblasts (Figure 5C, middle section), additionally supporting a functional role of PMAT in the feto-placental endothelium. We also analyzed expression of monoamine oxidase (MAOA), a rate-limiting enzyme in the catabolism of 5-HT^37^. *MAOA* mRNAs (Figure 5C, right section) were abundant in both cell types, with 4.9-fold higher levels (*p*<0.01) in trophoblasts than feto-placental endothelial cells.

**Figure 3.**
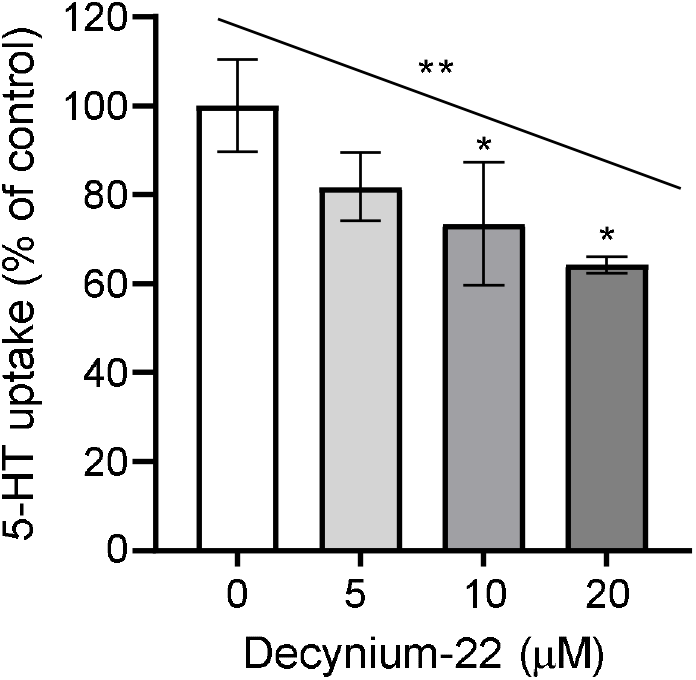
The effect of decynium-22, a common inhibitor of plasma membrane monoamine transporter and organic cation transporter 3, on 5-HT uptake into human primary feto-placental endothelial cells. Cells were incubated in the presence of ^3^H-5-HT (10^−4^ M) and increasing concentrations of decynium-22 for 10 min. Data are means ± SD of triplicates and shown is representative of two separate experiments. \**p*<0.05, compared to vehicle without drug (Holm-Sidak’s multiple comparison test), \*\**p*<0.01 for linear trend between column means (test for linear trend), both following one-way ANOVA (F_3,6_=6.687, *p*=0.024).

**Figure 4.**
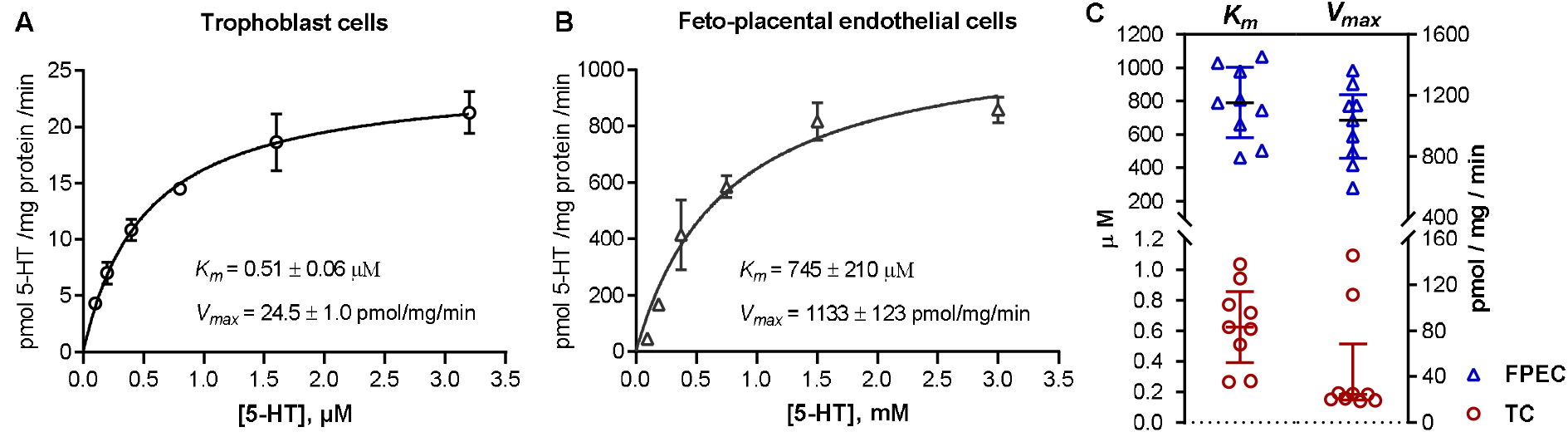
Kinetics of 5-HT uptake into human primary placental cells. Human primary (**A**) trophoblast cells and (**B**) feto-placental endothelial cells were incubated in the presence of six increasing ^3^H-5-HT concentrations ranging between (**A**) 0.1 and 3.2 μM, and (**B**) 94 and 3000 μM, for 2 or 8 min, respectively. Specific uptake was calculated as the difference between transport at 37°C and on ice. The initial rates of specific uptake were plotted against substrate concentration and fitted into Michaelis-Menten kinetics model. Shown are data (means ± SD, n=3) from one out of nine subjects (placentas) analyzed. (**C**) Best-fit values of Michaelis affinity constant (*Km*) and maximal transport velocity (*Vmax*) obtained in trophoblast cells (TC) and fetoplacental endothelial cells (FPEC) isolated from different placentas (n=9 for each cell type). Lines represent medians and interquartile range. Values of *Km* and *Vmax* were significantly different between trophoblasts and feto-placental endothelial cells (*p*<0.0001 in both cases; t-test and Mann Whitney test, respectively). For numerical results see Supplementary Table S1.

**Figure 5.**
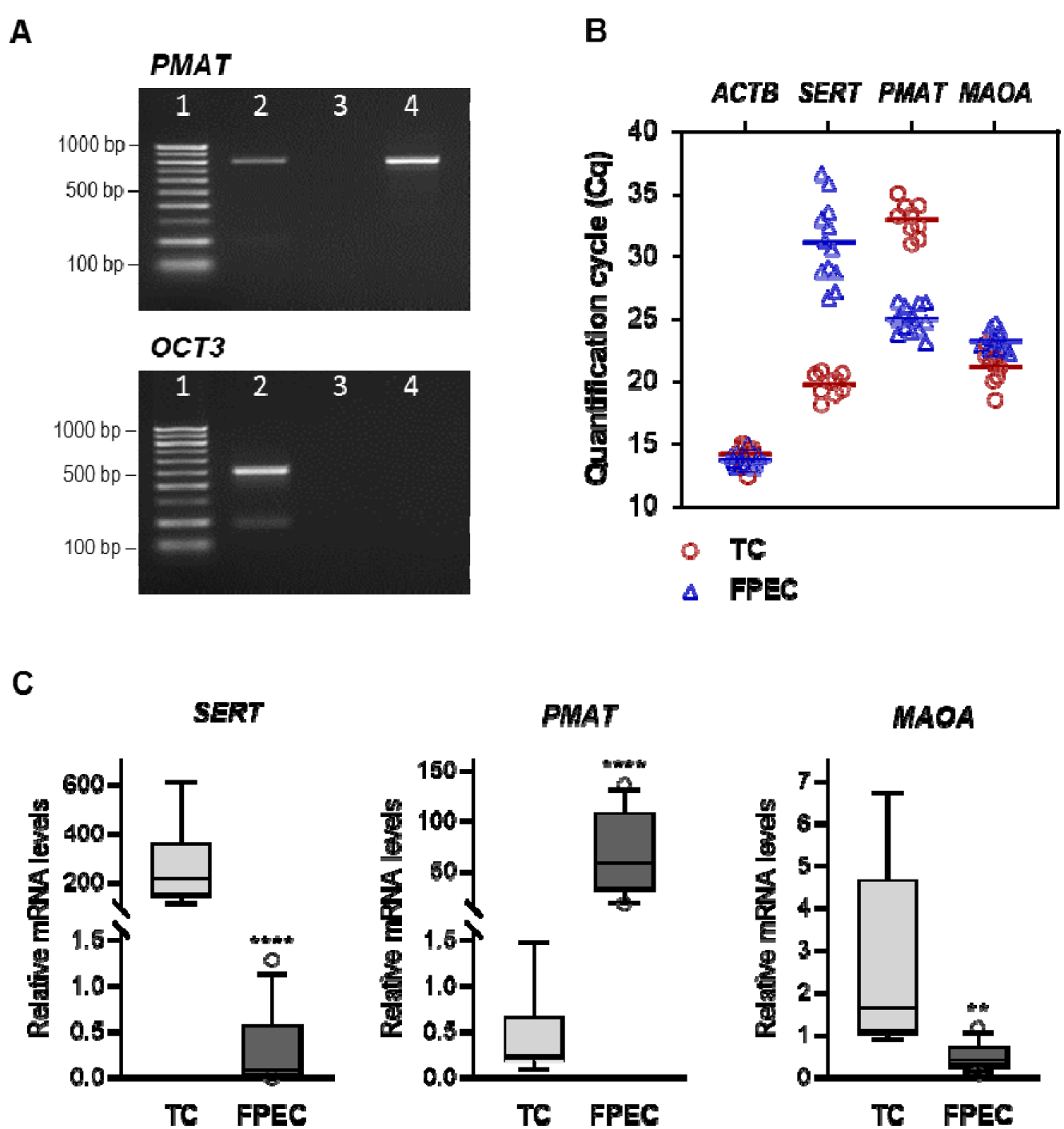
Expression of 5-HT regulating genes in primary placental cells. (**A**) Human primary feto-placental endothelial cells express plasma membrane monoamine transporter (*PMAT*) but not organic cation transporter 3 (*OCT3*) mRNA. Shown are products obtained by end-point RT-PCR analysis. The expected size of *PMAT* and *OCT3* specific amplicons corresponds to 762 and 504 bp, respectively^30^. Lane 1 - 100 bp DNA ladder, lane 2 - positive control (total placental tissue), lane 3 - control reaction lacking reverse transcriptase, lane 4 - feto-placental endothelial cells. (**B**) Quantification cycle (Cq) values obtained by RT-qPCR analysis of actin beta (*ACTB*), serotonin transporter (*SERT), PMAT* and monoaminoxidase A (*MAOA)* in human primary trophoblast cells (TC, n=9) and feto-placental endothelial cells (FPEC, n=12). Each dot is a mean of qPCR triplicates. (**C**) Relative expression levels of *SERT, PMAT* and *MAOA* mRNAs normalized to *ACTB* mRNA. Data are presented as boxplots with whiskers denoting the 10th and 90th percentiles (n= 9 and 12 for TC and FPEC, respectively). ^**^*p*<0.01, ^****^*p*<0.0001 (t-test or Mann-Whitney test, as appropriate).

### Effects of various psychotropic drugs on 5-HT uptake in human primary trophoblasts

Since SERT is a main target of many widespread medications and substances of abuse, we additionally tested potency of some of these drugs to inhibit 5-HT uptake into human primary trophoblasts. 5-HT uptake into trophoblasts was decreased in the presence of submicromolar (10^−7^ M) concentrations of all tested antidepressants (Figure 6A), namely, citalopram (by 91.8%), paroxetine (by 89.4%), fluoxetine (by 44.4%), fluvoxamine (by 53.0%), clomipramine (by 71.4%) and imipramine (by 32.3%). The SERT-interacting psychoactive stimulant 3,4-methylenedioxy-methamphetamine (MDMA, “Ecstasy”) also inhibited transport of 5-HT into trophoblasts, although with a lower potency - specifically, MDMA exhibited IC_50_ value in 10^−6^ M range, while those of citalopram and fluoxetine were in the range of only 10^−9^ and 10^−8^ M, respectively (Figure 6B).

**Figure 6.**
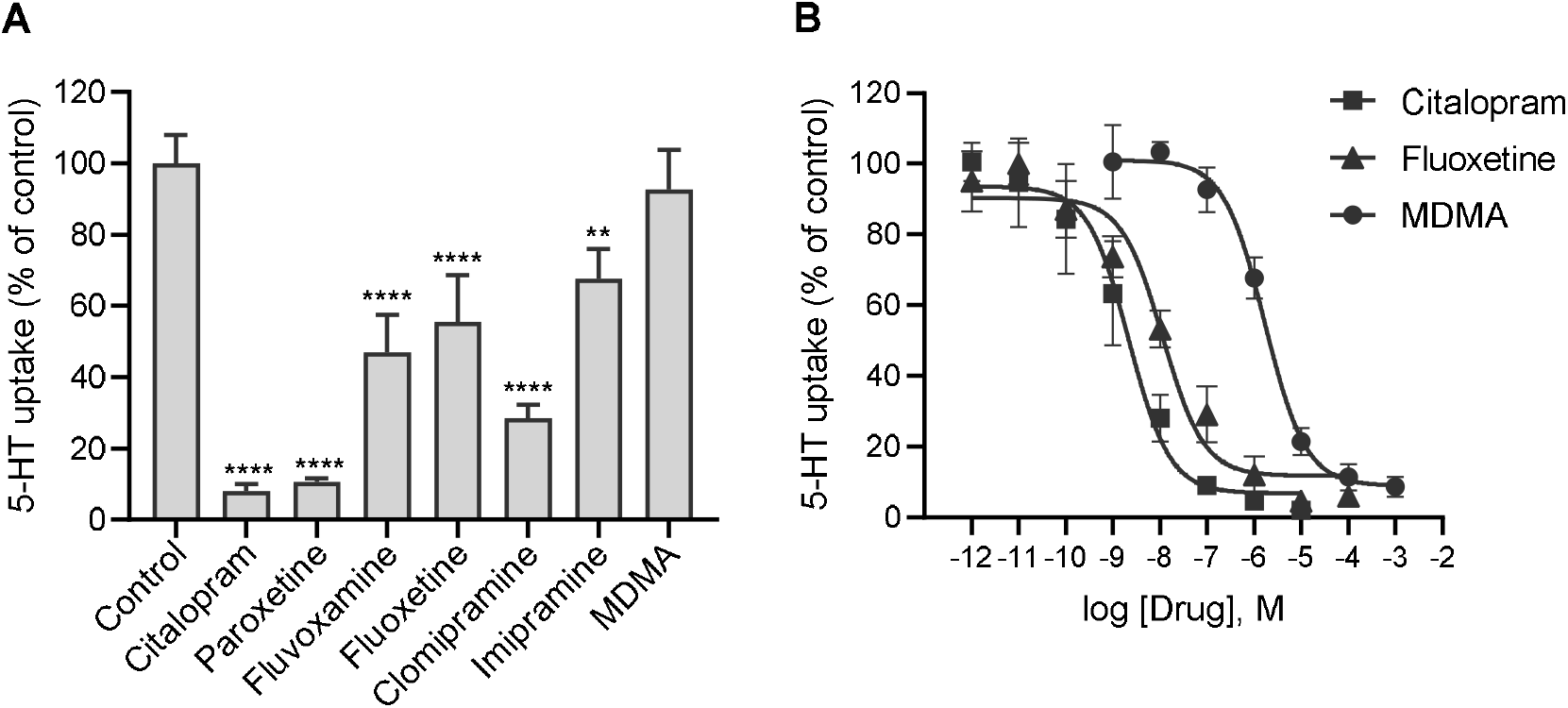
Potency of various psychotropic drugs to inhibit 5-HT uptake into human primary trophoblasts. Cells were incubated for 10 min in the presence of ^3^H-5-HT (10^−7^ M) and (**A**) 10^−7^ M or (**B**) increasing concentrations of the indicated drugs. Values are expressed as a percentage of the control (vehicle without the drug). All data are means ± SEM of three separate experiments, each performed in triplicate. Best fit half-maximal inhibitory concentrations (IC_50_) of citalopram, fluoxetine and MDMA amounted to 2, 11 and 1678 nM, with 95% confidence interval (CI) 1-7, 4-33 and 1045-2745 nM, respectively. \*\**p<0.01,* \*\*\*\**p<0.0001* for comparison to control (Dunnett’s test following one-way ANOVA, F_7,16_=52.34, *p*<0.0001). MDMA - 3,4-methylenedioxy-methamphetamine.

### 5-HT uptake in cord blood platelets

To examine potential further mechanisms contributing to uptake of 5-HT from the fetal circulation, we investigated 5-HT transport into platelets isolated from cord blood samples. Cord blood platelets showed efficient, time- and temperature-dependent 5-HT uptake, with initial rates of specific 5-HT transport saturable over the high-affinity range of 5-HT concentrations (0.15 to 2.0 μM; Figure 7A). *Km* values (0.654 ± 0.181 μM, n=9; Figure 7B) were in all subjects typical for a high-affinity uptake^26^ and similar to those found in trophoblasts (Figure 4C and Supplementary Table S1) and earlier in adult platelets^38,39^. *Vmax* values expressed per platelet number were also comparable between cord blood (109 ± 35 pmol / 10^8^ platelets /min) and adult (142 ± 25 pmol / 10^8^ platelets / min) ^38,39^ platelets. When expressed per total cellular protein, *Vmax* values were on average 20-fold higher (*p*<0.0001, Mann-Whitney test) in cord blood platelets (Figure 7B) than trophoblasts (Figure 4C; see also Supplementary Table S1). This is in line with platelets being among the few cell types with the highest amount of SERT protein in the adult human body^40^.

**Figure 7.**
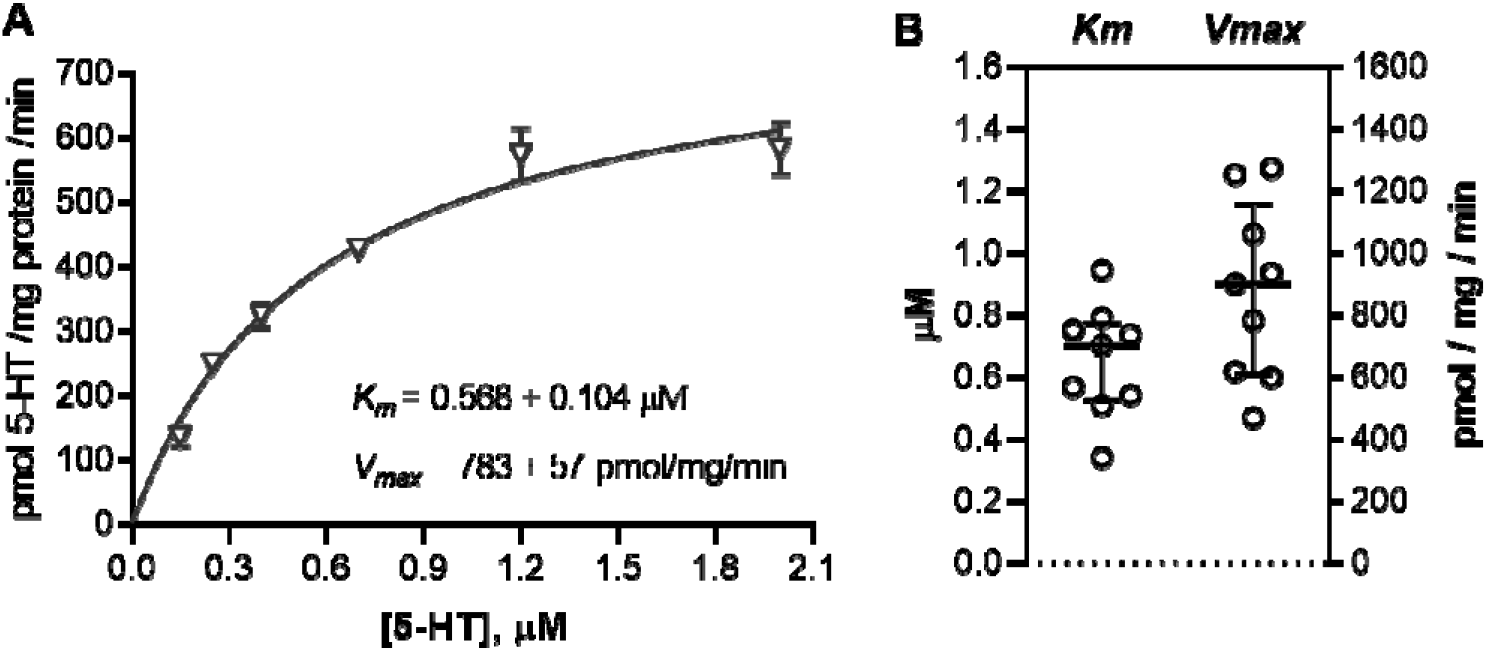
Uptake of 5-HT into human cord blood platelets. (**A**) The initial rates of specific 5-HT uptake plotted against substrate concentration and fitted into Michaelis-Menten kinetics model. Platelets were isolated from cord blood samples and incubated for 1 min in the presence of six ^14^C-5-HT concentrations (0.15 to 2.00 μM). Specific transport was determined as the difference between transport at 37°C and at around 4°C (ice bath). Shown are data (means ± SD) from one out of nine subjects analyzed. (**B**) Best-fit values of Michaelis affinity constant (*Km*) and maximal transport velocity (*Vmax*) in cord blood platelets isolated from different subjects (n=9). Lines represent medians and interquartile range. For numerical results see Supplementary Table S1.

## DISCUSSION

Here we investigated 5-HT uptake mechanisms in human primary placental cells and cord blood platelets, all isolated directly after birth. To determine uptake kinetics, we carried out comprehensive and rigorous initial rate studies using radiotracer assay. The initial rate period was experimentally determined for each cell type by time-dependent experiments. Specific (carrier-mediated) uptake was computed as a difference between total (at 37°C) and non-specific (at around 4°C) uptake. Initial rates of specific 5-HT uptake were measured at multiple substrate concentrations, covering a range typical for both high-affinity (uptake-1) and low-affinity (uptake-2) transport systems. This allowed estimation of kinetic parameters *Km* (approximating affinity of a carrier for 5-HT) and *Vmax* (describing maximal uptake velocity attained when a carrier is fully saturated). Additionally, we studied pharmacological properties of 5-HT uptake and examined mRNA levels of 5-HT carriers and 5-HT catabolizing enzyme in primary placental cells.

Our results show that human primary trophoblasts express *SERT* mRNA and take up 5-HT with a high-affinity transport kinetics characteristic of SERT-mediated process. *Vmax* values of 5-HT uptake in primary trophoblasts (18 - 146 pmol /mg /min, n=9) were similar to those reported for plasma membrane vesicles isolated from human native placentas (25.6 - 270 pmol /mg /min)^32,41–44^, but considerably higher than those reported for trophoblast-like cell line JAR (0.88 - 1.58 pmol /mg /min)^33,44–46^. These findings suggest a loss of functional SERT protein in commonly used JAR cells. They also indicate primary human trophoblasts as a credible physiological model for studying various regulatory features of placental 5-HT uptake from the maternal circulation in an intact cellular system.

Epidemiological data show that SERT-targeting antidepressant medications are increasingly used in pregnancy^47,48^. Therefore, we have evaluated the ability of these drugs to affect 5-HT uptake in human primary trophoblasts, a prime cell type on which antidepressants administered to the mother would act in the human feto-placental unit. We have found that 5-HT uptake into human primary trophoblasts was inhibited by nanomolar concentrations of common tricyclic (imipramine, clomipramine) and SSRI (citalopram, paroxetine, fluoxetine, fluvoxamine) antidepressants, with citalopram and paroxetine demonstrating the highest inhibitory effects. In addition, 5-HT uptake in trophoblasts was antagonized by the SERT-interacting recreational psychostimulant MDMA, the use of which also appears to increase among pregnant women^49^. It has been shown that both antidepressants and MDMA may actively traverse the placenta and affect the developing fetal organs^50–52^. Our results in human primary trophoblasts together with earlier findings in human placental explants^14^ and plasma membrane vesicles^32,41,42^ highlight that placenta itself, specifically, its mother-facing side, may be a target of these drugs. Potential downstream consequences of disturbed placental 5-HT homeostasis induced by inhibition of 5-HT uptake in mother-facing trophoblast cells warrant further investigation.

Mechanisms involved in uptake of 5-HT from the fetal circulation have only recently gained attention^34^. Here we have for the first time demonstrated functional 5-HT uptake system at term of human pregnancy in two cell types that are in direct contact with the fetal blood plasma. First, we uncovered a low-affinity (most likely PMAT-mediated) 5-HT uptake activity in feto-placental endothelial cells. Second, we demonstrated a high-affinity 5-HT uptake activity in cord blood platelets, with kinetic parameters resembling to that of SERT-mediated 5-HT transport into adult platelets^38,39^. Additionally, we identified expression of MAOA, a 5-HT catabolizing enzyme with a high affinity for 5-HT, in feto-placental endothelial cells. This suggests that feto-placental endothelial cells can catabolize 5-HT to an inactive metabolite. In platelets, 5-HT that has been taken up may be either sequestered in dense granules or catabolized by MAOB^53^. Taken together, these findings suggest that both feto-placental endothelial cells and fetal platelets have systems in place to actively participate in the uptake and deactivation of 5-HT from the fetal circulation.

The observed *Km* values of 5-HT uptake in feto-placental endothelial cells (782 ± 218 μM) were about three orders of magnitude higher than that in cord blood platelets (0.654 ± 018 μM), suggesting that the two uptake systems operate at fundamentally different substrate concentrations. We speculate that feto-placental endothelial cells and fetal platelets cooperatively work to meet requirements for maintaining optimal *in vivo* levels of extracellular 5-HT in different situations (Figure 8). Under basal conditions, when fetal plasma 5-HT concentrations are in the low nanomolar range^54^, uptake of 5-HT is mediated principally by the high-affinity system of fetal platelets. When fetal plasma 5-HT concentrations increase to levels at which the high-affinity system of platelets is fully saturated, the low-affinity system of feto-placental endothelial cells, which saturates at much higher concentrations, takes part in helping to bring extracellular 5-HT to low basal levels. A further pathway of 5-HT sequestration in the human placenta, operating especially at high 5-HT levels in the fetal plasma, may imply 5-HT diffusion through paracellular routes between feto-placental endothelial cells and placental stroma to reach the syncytiotrophoblast basal membrane. There it might serve as substrate of the low-affinity, OCT3-mediated uptake system^34^. Extracellular 5-HT concentrations may fluctuate locally / transiently towards high levels as a consequence of 5-HT release from fetal platelets or other cells, for endocrine, paracrine or autocrine signaling purposes. Other situations in which the low-affinity 5-HT uptake systems present in feto-placental endothelial cells and syncytiotrophoblast basal membrane may be important are when the SERT-mediated uptake in fetal platelets is inhibited pharmacologically or is compromised due to genetic or pathology-related reasons. Indeed, maternal treatment with SSRI antidepressants has been associated with decreased 5-HT levels in cord blood platelets^55^, suggesting inhibited uptake of 5-HT into fetal platelets. There is evidence that the 5-HT clearance system in the brain also combines high-affinity and low-affinity transporters^56^. Interplay between different 5-HT uptake systems in the feto-placental unit warrants further investigation.

**Figure 8.**
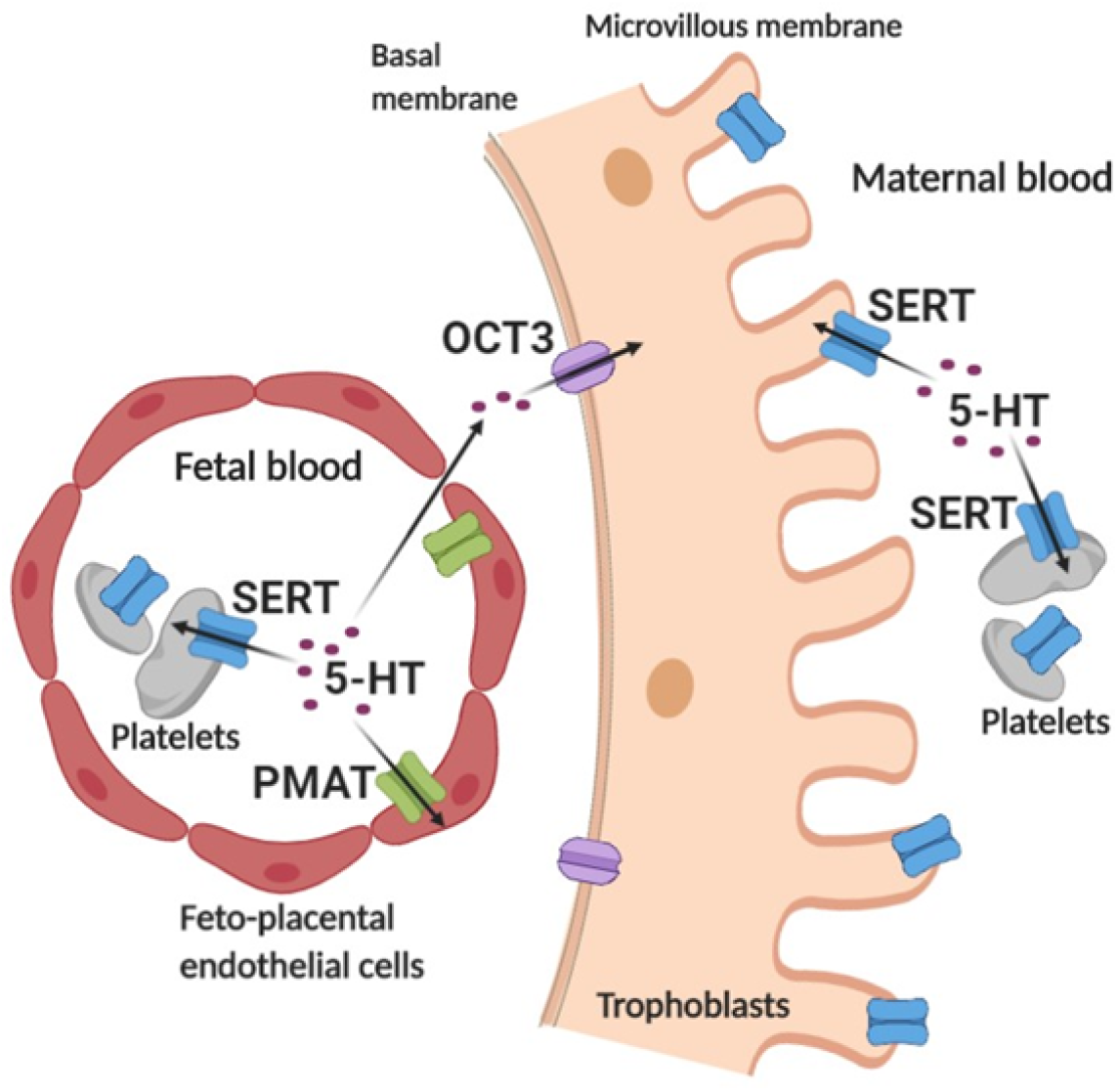
Multiple systems for cellular uptake of serotonin from placental extracellular space at the end of human pregnancy. Serotonin (5-HT) in fetal plasma is taken up via high-affinity serotonin transporter (SERT) into fetal platelets and via low-affinity plasma membrane monoamine transporter (PMAT) into fetoplacental endothelial cells. In addition, fetal 5-HT may diffuse through paracellular routes between fetoplacental endothelial cells and placental stroma to the basal side of trophoblasts where it serves as a substrate of a low-affinity organic cation transporter 3 (OCT3)^34^. 5-HT in maternal plasma is sequestrated via SERT located on both maternal platelets and apical side of trophoblasts. Created with BioRender.com.

We have to acknowledge the weaknesses of our study. First, different types of cells analyzed were not obtained from the same subject or placenta as a consequence of technical constraints. We used a larger number of different donors for kinetic and gene expression analyses to account for this issue. Further, primary placental cells were not maintained under physiological oxygen conditions, which would better mimic the physiological environment. Oxygen level was reported to modulate SERT activity in certain human cells^57^ and thus may have influenced the results. Third, it should be noted that primary trophoblasts obtained from different donors showed an extensive inter-individual variability in *Vmax* (up to 8.1-fold) and *Km* (up to 3.9-fold) values. Considerable inter-individual differences were present also in *Vmax* and *Km* values in feto-placental-endothelial cells (both by up to 2.3-fold) and fetal platelets (by up to 2.7- and 2.8-fold, respectively). This might be a consequence of the influence of various fetal and/or maternal factors that were not controlled for in the present study. Indeed, fetal sex^34^ and genotype^58^ were reported to influence 5-HT uptake in the plasma membrane vesicles isolated from rat and human placentas, respectively. Various other gestational factors such as maternal obesity and glucose tolerance^17,36,59,60^ as well as maternal genotype, diet, stress and immune activation^22^ have been also reported to influence placental or fetal 5-HT homeostasis. Understanding the regulatory influences of various factors on the 5-HT uptake systems in placenta and platelets, which was beyond the scope of the present study, warrants further investigation in studies employing full kinetic measurements.

In summary, we comprehensively characterized kinetic and pharmacological aspects of 5-HT uptake into human primary trophoblasts facing maternal blood at the term. Besides establishing these cells as a valuable tool for studying different regulatory features of placental 5-HT uptake from maternal circulation, the results emphasize sensitivity of placental 5-HT transport to inhibitory effects of various psychotropic drugs. Further, we demonstrated the presence of a functional 5-HT uptake system, with low substrate affinity, in human feto-placental endothelial cells. These cells on the fetal side of the feto-placental unit are in direct contact with fetal blood and, hence, take up fetal 5-HT. Finally, we have shown that human fetal platelets express a functional high-affinity 5-HT uptake system resembling that in adult platelets. The present identification of multiple membrane transport systems for uptake of extracellular 5-HT at the human maternal-fetal interface (Figure 8) advances our understanding of mechanisms that govern 5-HT homeostasis during human fetal development. Since cellular 5-HT uptake is central to regulating local levels of 5-HT nearby its molecular targets, these results may open new opportunities for the modulation of disorders associated with developmental changes in 5-HT signaling.

## METHODS

### Materials

Tritium-labelled serotonin creatinine sulphate (^3^H-serotonin; 28.3 Ci mmol^−1^, 41.3 Ci mmol^−1^) and radiocarbon-labelled serotonin binoxalate (^14^C-serotonin; 54.0 mCi mmol^−1^) were purchased from Perkin Elmer (Waltham, Massachusetts, USA). Serotonin creatinine sulphate, imipramine hydrochloride, citalopram hydrobromide, 3,4-methylenedioxymethamphetamine hydrochloride (MDMA, ‘Ecstasy’), decynium 22 (D-22), L-ascorbic acid and Hanks’ Balanced Salts Solution (HBSS) were obtained from Sigma-Aldrich (St. Louis, Missouri, USA). Pargyline hydrochloride was from Cayman Chemical (Ann Arbor, Michigan, USA). Other drugs were generous gifts from the manufacturers: fluoxetine hydrochloride from Eli Lilly and Company (USA), paroxetine hydrochloride from GlaxoSmithKline (England), fluvoxamine maleate from Solvay (Netherlands) and Clomipramine from BASF Schweiz AG (Switzerland).

### Isolation and culture of human primary placental cells

The study was performed in accordance with the protocol approved by the Medical University of Graz, Graz, Austria. An informed written consent was obtained from all women who donated placentas for cell isolation. All placentas were from singleton term (37-42 weeks of gestation) pregnancies. Exclusion criteria were any known fetal or newborn anomalies. Additional information on participants is provided in Supplementary Table S2. Primary trophoblasts^61^ and feto-placental endothelial cells^62^ were isolated from villous tissue, as described earlier. Representative cell isolations were tested for identity and purity by immunocytochemical staining of specific cell markers^61,62^. Cells were cultured on gelatine (1%)-coated plates, trophoblasts in Dulbecco’s Modified Eagle’s Medium (DMEM; Gibco, Paisley, UK) and feto-placental endothelial cells in Endothelial Cell Basal Medium (EBM; Lonza, Verviers, Belgium) or Endothelial Cell Growth Medium MV (PromoCell, Heidelberg, Germany), all supplemented with 10% fetal bovine serum and 1% antibiotic. Cultures were maintained at 37°C, 5% CO_2_ and 21% O_2_.

### 5-HT uptake in human primary placental cells

Cells were seeded onto 24-well plates and allowed to reach confluency (usually for 1 to 2 days). Uptake was initiated, following removal of media and rinsing of cells with HBSS, by the addition of pre-warmed (37°C) or pre-cooled (4°C) uptake buffer (0.2 mL/well). The uptake buffer was HBSS containing a mixture of ^3^H-labeled and unlabeled 5-HT totaling the required final 5-HT concentration, as well as ascorbic acid (100 μM) and pargyline (10 μM) to prevent 5-HT oxidation and enzymatic degradation, respectively. Reactions were carried out at 37°C or on ice for a specified amount of time and were terminated by removal of the uptake buffer and addition of ice-cold HBSS (1 mL/well). Cells were then thoroughly washed with ice-cold HBSS and lysed in 0.3 N NaOH (0.4 mL/well). Radioactivity in cell lysates was quantified on Tri-Carb 2100TR Liquid Scintillation Counter, using Ultima Gold liquid scintillation cocktail (both from Perkin Elmer, Waltham, Massachusetts, USA). Total protein concentration in cell lysates was determined using Qubit™ Protein Assay Kit (Thermo Fisher Scientific, Waltham, Massachusetts, USA). Specific (carrier-mediated) uptake was calculated as the difference between total (at 37°C) and non-specific (on ice) uptake. All assays were performed in triplicates. In initial rate studies, 5-HT uptake was measured within experimentally determined linear time range, at six increasing 5-HT concentrations covering a range typical for high-affinity (0.1, 0.2, 0.4, 0.8. 1.6 and 3.2 μM) or low-affinity (94, 188, 375, 1500 and 3000 μM) uptake mechanism. The initial rates of specific 5-HT uptake were fitted into Michaelis-Menten kinetics model using GraphPad Prism software. Best-fit values of Michaelis affinity constant (*Km*) and maximal transport velocity (*Vmax*) were estimated by nonlinear least-squares regression analysis.

### Pharmacological studies

Stock solutions of MDMA (10^−1^ M) and D-22 (10^−2^ M) were prepared in DMSO, and of all other drugs (10^−2^ M) in HBSS. Following removal of media and rinsing with HBSS, cells were pre-incubated for 10 min in the presence of either drugs or vehicle (control) and then incubated for 10 min in the presence of ^3^H-5-HT and either drugs or vehicle, respectively. Transport on ice was subtracted from that at 37°C and specific uptake in the presence of each drug was expressed as percentage of the control (vehicle). There were no differences in 5-HT uptake between controls prepared with DMSO and HBSS at a concentration equivalent to that in samples with drugs. Half maximal inhibitory concentration values (IC_50_) with 95% confidence intervals (CI) were determined by GraphPad Prism software, using nonlinear least-squares regression analysis.

### Gene expression analyses

Cells used for gene expression analyses were seeded on 6-well plates and harvested at confluency. Placental tissue used as a positive control in qualitative end-point reverse transcription (RT) - polymerase chain reaction (PCR) was obtained as described earlier^60^. Total RNA was extracted by RNeasy Plus Mini Kit (Qiagen, Hilden, Germany), according to the manufacturer’s protocol with optional on-column DNA digestion step. Concentration and purity of RNA was determined spectrophotometrically (NanoDrop, Witec AG, Littau, Germany). RNA integrity was assessed by 1% agarose gel electrophoresis. cDNA was synthesized from uniform amounts of RNA, using iScript cDNA Synthesis Kit (Bio-Rad) according to the manufacturer’s protocol. Control reactions lacking reverse transcriptase (no-RT) were prepared to check for genomic DNA contamination. Sequences of primers used in end-point PCR and quantitative real-time PCR (qPCR) are listed in Supplementary Table S3. End-point PCR assays were performed with AllTaq Master Mix (Qiagen, Hilden, Germany), following manufacturer’s recommendations, and the obtained products were separated on 2% agarose gels stained with Midori Green Advance DNA Stain (Nippon Genetics, Germany). qPCR assays were performed on 7300 Real Time PCR System using Syber Green Master Mix (both from Applied Biosystems, Waltham, Massachusetts, USA), according to manufacturers’ recommendation. The specificity of qPCR products was verified by agarose gel electrophoresis and routinely by melting curve analysis. All qPCR assays were run in triplicates. qPCR efficiencies of genes of interest were about equal to that of *ACTB* used as a reference gene (the slope of log input amount versus ΔC_q_ < 0.1 in all cases) and relative expression levels were calculated using the comparative C_q_ (ΔΔC_q_) method^63^.

### Preparation of cord blood platelet rich plasma

The study protocol was approved by the Ethics Committee of the Clinical Hospital Centre Zagreb and Bioethics Committee of the Ruder Bośkoviċ Institute, Zagreb, Croatia. All methods were performed in accordance with relevant guidelines and regulations. Cord blood was obtained from singleton neonates born at term (37-42 weeks of gestation) by elective Caesarean section (Supplementary Table S2), following a signed informed consent provided by their mothers. Cord blood samples were collected via umbilical venipuncture after baby birth and before delivery of the placenta, and were immediately mixed with acid citrate dextrose (ACD) anticoagulant in a 1 : 5 (ACD : blood) ratio. Platelet rich plasma (PRP) was isolated by centrifugation of anti-coagulated blood for 2 min at 1100 g. Total platelet protein levels were determined by Bradford’s assay following centrifugation of PRP samples. Platelet number and volume were determined in whole blood and PRP samples using DxH 500 hematology analyzer (Beckman Coulter). Platelet count (mean ± SD, n=9) were 242 ± 112 x 10^6^ /mL in whole cord blood and 388 ± 92 x 10^6^ /mL in PRP samples. Mean platelet volumes (mean ± SD, n=9) in whole cord blood (7.8 ± 0.7 fL) and PRP (7.4 ± 0.62 fL) samples were highly correlated (Spearmen’s correlation coefficient = 0.94, *p*=0.001, n=9), demonstrating that population of platelets isolated in PRP represented well platelets in the whole cord blood samples.

### 5-HT uptake in cord blood platelets

Uptake of 5-HT in cord blood platelets was measured within two hours after sampling, according to a slightly modified protocol used in our previous studies in adults^39,61^. In brief, PRP samples (60 μL) were mixed with CaCl_2_-free Krebs-Ringer phosphate buffer (KRB; 840 μL) and pre-incubated for 10 min in a shaking bath at 37°C. Uptake was initiated by the addition of KRB (100 μL) containing ^14^C-5-HT. Final concentrations of ^14^C-5-HT in the reaction mixture amounted to 0.15, 0.25, 0.40, 0.70, 1.20 and 2.00 μM. Reactions were carried out at 37°C for 60 s and were terminated by rapid cooling (addition of ice-cold saline) and immediate vacuum filtration over a glass microfiber filters (Whatman GF/C, GE Healthcare, Illinois, USA). The radioactivity retained on the filters after thorough rinsing was quantified on Tri-Carb 2810TR Liquid Scintillation Analyzer, using Ultima Gold MV liquid scintillation cocktail (both from Perkin Elmer, Waltham, Massachusetts, USA). Non-specific uptake was measured by the same procedure, but at around 4°C (ice bath). All assays were performed in duplicate. Specific (carrier-mediated) uptake, calculated as a difference between total (at 37°C) and nonspecific (ice bath) uptake, was expressed per platelet number (for comparison with published data for adult platelets) and per total platelet protein (for comparison with results in human primary trophoblasts and feto-placental endothelial cells). Values of *Km* and *Vmax* were determined as described for primary placental cells.

### Statistical analysis

Statistical analyses were conducted using GraphPad Prism version 8 (GraphPad Software, LLC, San Diego, CA, USA). Data distribution was tested by D’Agostino-Pearson omnibus normality test. Normally distributed data were analyzed by Student’s t-test, one-way analysis of variance (ANOVA) or two-way ANOVA. Not normally distributed data were analyzed by Mann-Whitney test. Statistical tests applied are specified in Figure legends. The level of significance was set at 0.05.

## DATA AVAILABILITY

The datasets generated during and/or analyzed during the current study are available from the corresponding author on reasonable request.

## ACKNOWLEDGMENTS

The authors thank Susanne Kopp and Renate Michlmaier for their assistance in isolating primary placental cells. The authors also thank participants who donated tissues for the study. This work was supported by the Croatian Science Foundation (grant number: IP-2018-01-6547).

## AUTHORS’ CONTRIBUTIONS

JŠ and UP conceived the study and supervised the experiments. JŠ, UP, CW, LCŠ and GD designed the experiments and interpreted the results. PB, MK, MH, MG, MP, IB and JŠ carried out the experiments and analyzed the data. JŠ wrote and GD, LCS, IB and CW critically revised the manuscript. UP deceased during the preparation of the manuscript; other authors read and approved the final version of the manuscript.

## SUPPLEMENTARY MATERIAL

**Supplementary Table S1.** Kinetic parameters of 5-HT uptake into human primary trophoblasts, fetoplacental endothelial cells and cord blood platelets. Shown are means ± SD or, where appropriate, medians with interquartile range (IQR, in parenthesis). Data for trophoblasts and feto-placental endothelial cells are derived from Figures 4C and for cord blood platelets from Figure 7B. *Km* - Michaelis affinity constant, *Vmax* - maximal transport velocity, N - number of participants analyzed.

| Cells | N | *K<sub>m</sub>* (μM) | *V<sub>max</sub>* (pmol /mg protein /min) |
| --- | --- | --- | --- |
| Trophoblasts | 9 | 0.640 ± 0.266 | 24.5 (48.7) |
| Feto-placental endothelial cells | 9 | 782 ± 218 | 1005 ± 251 |
| Cord blood platelets | 9 | 0.654 ± 0.181 | 878 ± 286 |

**Supplementary Table S2.**
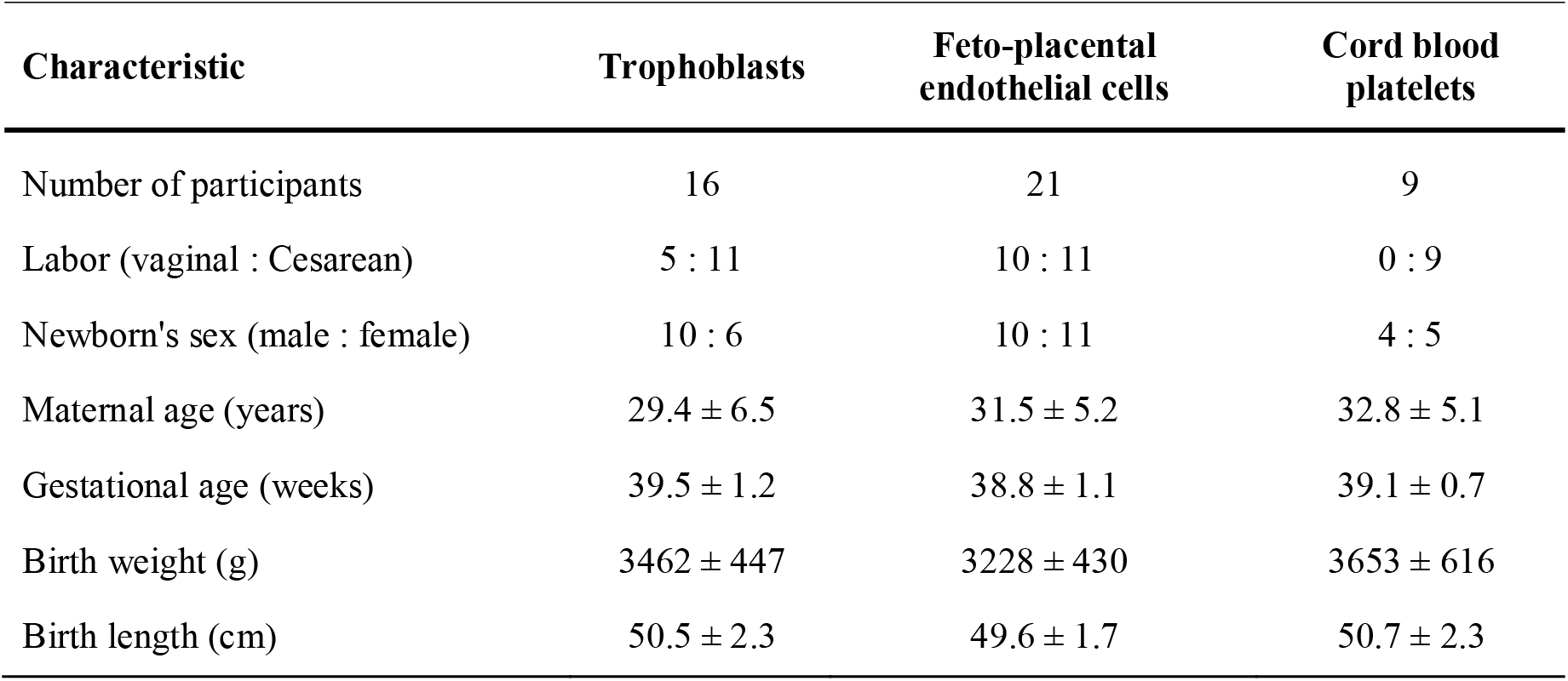
Characteristics of participants who donated tissues for isolation of trophoblasts, feto-placental endothelial cells and cord blood platelets used in pharmacological and kinetic 5-HT uptake studies and gene expression analyses. Continuous variables are given as mean ± SD.

**Supplementary Table S3.**
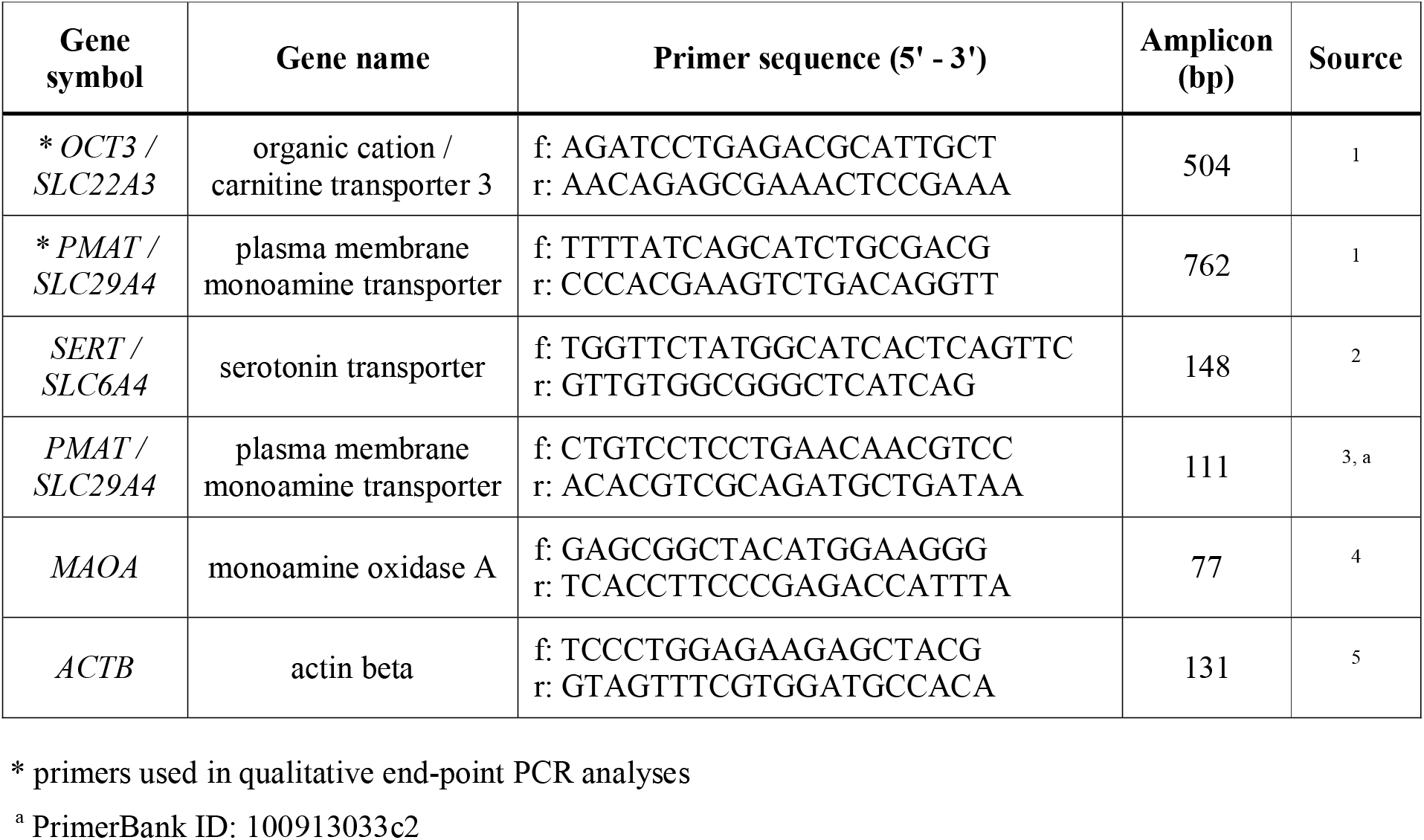
Sequences of gene-specific primers used in qualitative end-point PCR and quantitative real-time PCR analyses. f - forward, r - reverse.

## Supplementary Figures

**Supplementary Figure S1.**
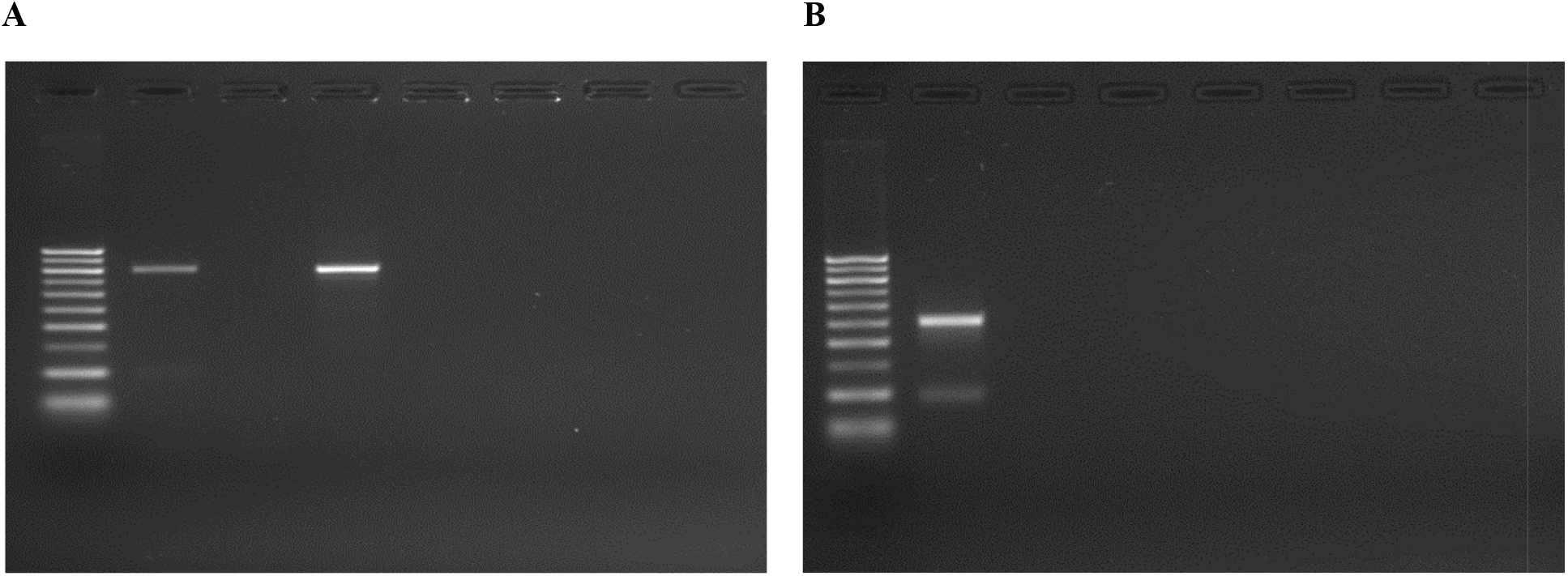
Uncropped and unprocessed agarose gel electrophoresis images of (**A**) *PMAT* and (**B**) *OCT3* end-point RT-PCR experiments shown in the main text as Figure 5A, upper and lower section, respectively. Order of samples on both images: lane 1 - 100 bp ladder, lane 2 - total placental tissue (positive control), lane 3 - control reaction lacking reverse transcriptase (with RNA from total placental tissue), lane 4 - feto-placental endothelial cells, lane 5 - control reaction lacking reverse transcriptase (with RNA from fetoplacental endothelial cells).

## REFERENCES

1. Berger, M., Gray, J. A. & Roth, B. L. The expanded biology of serotonin. Annu. Rev. Med. 60, 355–366 (2009).

2. Spohn, S. N. & Mawe, G. M. Non-conventional features of peripheral serotonin signalling-the gut and beyond. Nat. Rev. Gastroenterol. Hepatol. 14, 412–420 (2017).

3. Gaspar, P., Cases, O. & Maroteaux, L. The developmental role of serotonin: News from mouse molecular genetics. Nat. Rev. Neurosci. 4, 1002–1012 (2003).

4. Rosenfeld, C. S. Placental serotonin signaling, pregnancy outcomes, and regulation of fetal brain development†. Biol. Reprod. 102, 532–538 (2020).

5. Sonier, B., Lavigne, C., Arseneault, M., Ouellette, R. & Vaillancourt, C. Expression of the 5-HT2A serotoninergic receptor in human placenta and choriocarcinoma cells: mitogenic implications of serotonin. Placenta. 26, 484–490 (2005).

6. Oufkir, T., Arseneault, M., Sanderson, J. T. & Vaillancourt, C. The 5-HT2A serotonin receptor enhances cell viability, affects cell cycle progression and activates MEK-ERK1/2 and JAK2-STAT3 signalling pathways in human choriocarcinoma cell lines. Placenta. 31, 439–447 (2010).

7. Oufkir, T. & Vaillancourt, C. Phosphorylation of JAK2 by serotonin 5-HT2A receptor activates both STAT3 and ERK1/2 pathways and increases growth of JEG-3 human placental choriocarcinoma cell. Placenta. 32, 1033–1040 (2011).

8. Klempan, T., Hudon-Thibeault, A. A., Oufkir, T., Vaillancourt, C. & Sanderson, J. T. Stimulation of serotonergic 5-HT2A receptor signaling increases placental aromatase (CYP19) activity and expression in BeWo and JEG-3 human choriocarcinoma cells. Placenta. 32, 651–656 (2011).

9. Hadden, C. et al. Serotonin transporter protects the placental cells against apoptosis in caspase 3-independent pathway. J. Cell. Physiol. 232, 3520–3529 (2017).

10. Cruz, M. A., Gallardo, V., Miguel, P., Carrasco, G. & González, C. Serotonin-induced vasoconstriction is mediated by thromboxane release and action in the human fetal-placental circulation. Placenta. 18, 197–204 (1997).

11. Gonzalez, C., Cruz, M. A., Gallardo, V., Albornoz, J. & Bravo, I. Serotonin-induced vasoconstriction in human placental chorionic veins: Interaction with prostaglandin F2α. Gynecol. Obstet. Invest. 35, 86–90 (1993).

12. Côté, F. et al. Maternal serotonin is crucial for murine embryonic development. Proc. Natl. Acad. Sci. U.S.A. 104, 329–334 (2007).

13. Bonnin, A. et al. A transient placental source of serotonin for the fetal forebrain. Nature. 472, 347–350 (2011).

14. Kliman, H. J. et al. Pathway of maternal serotonin to the human embryo and fetus. Endocrinology. 159, 1609–1629 (2018).

15. Carrasco, G. et al. Transport and metabolism of serotonin in the human placenta from normal and severely pre-eclamptic pregnancies. Gynecol. Obstet. Invest. 49, 150–155 (2000).

16. Ranzil, S. et al. Disrupted placental serotonin synthetic pathway and increased placental serotonin: potential implications in the pathogenesis of human fetal growth restriction. Placenta. 84, 74–83 (2019).

17. Murthi, P. & Vaillancourt, C. RETRACTED: Placental serotonin systems in pregnancy metabolic complications associated with maternal obesity and gestational diabetes mellitus. Biochim. Biophys. Acta. Mol. Basis. Dis. 1866, 65391 (2020).

18. Bonnin, A. & Levitt, P. Fetal, maternal, and placental sources of serotonin and new implications for developmental programming of the brain. Neuroscience. 197, 1–7 (2011).

19. Sato, K. Placenta-derived hypo-serotonin situations in the developing forebrain cause autism. Med. Hypotheses. 80, 368–372 (2013).

20. Räikkönen, K. et al. Maternal depressive symptoms during pregnancy, placental expression of genes regulating glucocorticoid and serotonin function and infant regulatory behaviors. Psychol. Med. 45, 3217–3226 (2015).

21. Goeden, N. et al. Maternal inflammation disrupts fetal neurodevelopment via increased placental output of serotonin to the fetal brain. J. Neurosci. 36, 6041–6049 (2016).

22. Hanswijk, S. I. et al. Gestational factors throughout fetal neurodevelopment: the serotonin link. Int. J. Mol. Sci. 21, 5850 (2020).

23. Hoyer, D., Hannon, J. P. & Martin, G. R. Molecular, pharmacological and functional diversity of 5-HT receptors. Pharmacol. Biochem. Behav. 71, 533–554 (2002).

24. Muma, N. A. & Mi, Z. Serotonylation and transamidation of other monoamines. ACS Chem. Neurosci. 6, 961–969 (2015).

25. Farrelly, L. A. et al. Histone serotonylation is a permissive modification that enhances TFIID binding to H3K4me3. Nature. 567, 535–539 (2019).

26. Ramamoorthy, S. et al. Antidepressant- and cocaine-sensitive human serotonin transporter: molecular cloning, expression, and chromosomal localization. Proc. Natl. Acad. Sci. U.S.A. 90, 2542–2546 (1993).

27. Mercado, C. P. & Kilic, F. Molecular mechanisms of SERT in platelets: regulation of plasma serotonin levels. Mol. Interv. 10, 231–241 (2010).

28. Duan, H. & Wang, J. Selective transport of monoamine neurotransmitters by human plasma membrane monoamine transporter and organic cation transporter 3. J. Pharmacol. Exp. Ther. 335, 743–753 (2010).

29. Li, R. W. S. et al. Involvement of organic cation transporter-3 and plasma membrane monoamine transporter in serotonin uptake in human brain vascular smooth muscle cells. Front. Pharmacol. 4, 14 (2013).

30. Naganuma, F. et al. Predominant role of plasma membrane monoamine transporters in monoamine transport in 1321N1, a human astrocytoma-derived cell line. J. Neurochem. 129, 591–601 (2014).

31. Wang, J. The plasma membrane monoamine transporter (PMAT): structure, function, and role in organic cation disposition. Clin. Pharmacol. Ther. 100, 489–499 (2016).

32. Balkovetz, D. F., Tiruppathi, C., Leibach, F. H., Mahesh, V. B. & Ganapathy, V. Evidence for an imipramine-sensitive serotonin transporter in human placental brush-border membranes. J. Biol. Chem. 264, 2195–2198 (1989).

33. Cool, D. R., Leibach, F. H., Bhalla, V. K., Mahesh, V. B. & Ganapathy, V. Expression and cyclic AMP-dependent regulation of a high affinity serotonin transporter in the human placental choriocarcinoma cell line (JAR). J. Biol. Chem. 266, 15750–15757 (1991).

34. Karahoda, R. et al. Serotonin homeostasis in the materno-foetal interface at term: Role of transporters (SERT/SLC6A4 and OCT3/SLC22A3) and monoamine oxidase A (MAO-A) in uptake and degradation of serotonin by human and rat term placenta. Acta. Physiol. (Oxf.). 229, e13478 (2020).

35. Hyttel, J. Pharmacological characterization of selective serotonin reuptake inhibitors (SSRIs). Int. Clin. Psychopharmacol. 9, 19–26 (1994).

36. Viau, M., Lafond, J. & Vaillancourt, C. Expression of placental serotonin transporter and 5-HT2A receptor in normal and gestational diabetes mellitus pregnancies. Reprod. Biomed. Online. 19, 207–215 (2009).

37. Shih, J. C. Monoamine oxidase isoenzymes: genes, functions and targets for behavior and cancer therapy. J. Neural Transm. (Vienna). 125, 1553–1566 (2018).

38. Banović, M., Bordukalo-Nikśić, T., Balija, M., Cičm-Šam, L. & Jernej, B. Platelet serotonin transporter (5HTt): physiological influences on kinetic characteristics in a large human population. Platelets. 21, 429–438 (2010).

39. Balija, M. et al. Serotonin level and serotonin uptake in human platelets: a variable interrelation under marked physiological influences. Clin. Chim. Acta. 412, 299–304 (2011).

40. Beikmann, B. S., Tomlinson, I. D., Rosenthal, S. J. & Andrews, A. M. Serotonin uptake is largely mediated by platelets versus lymphocytes in peripheral blood cells. ACS Chem. Neurosci. 4, 161–170 (2013).

41. Cool, D. R., Leibach, F. H. & Ganapathy, V. High-affinity paroxetine binding to the human placental serotonin transporter. Am. J. Physiol. 259, C196–204 (1990).

42. Cool, D. R., Liebach, F. H. & Ganapathy, V. Interaction of fluoxetine with the human placental serotonin transporter. Biochem. Pharmacol. 40, 2161–2167 (1990).

43. Ramamoorthy, S., Cool, D. R., Leibach, F. H., Mahesh, V. B. & Ganapathy, V. Reconstitution of the human placental 5-hydroxytryptamine transporter in a catalytically active form after detergent solubilization. Biochem. J. 286, 89–95 (1992).

44. Prasad, P. D., Leibach, F. H., Mahesh, V. B. & Ganapathy, V. Human placenta as a target organ for cocaine action: Interaction of cocaine with the placental serotonin transporter. Placenta. 15, 267–278 (1994).

45. Jayanthi, L. D., Ramamoorthy, S., Mahesh, V. B., Leibach, F. H. & Ganapathy, V. Calmodulindependent regulation of the catalytic function of the human serotonin transporter in placental choriocarcinoma cells. J. Biol. Chem. 269, 14424–14429 (1994).

46. Decker, A. M. & Blough, B. E. Development of serotonin transporter reuptake inhibition assays using JAR cells. J. Pharmacol. Toxicol. Methods. 92, 52–56 (2018).

47. Andrade, S. E. et al. Use of antidepressant medications during pregnancy: a multisite study. Am. J. Obstet. Gynecol. 198, 194.e1–194.e5 (2008).

48. Bakker, M. K., Kölling, P., Van Den Berg, P. B., De Walle, H. E. K. & De Jong van den Berg, L. T. W. Increase in use of selective serotonin reuptake inhibitors in pregnancy during the last decade, a population-based cohort study from the Netherlands. Br. J. Clin. Pharmacol. 65, 600–606 (2008).

49. Smid, M. C., Metz, T. D. & Gordon, A. J. Stimulant use in pregnancy: an under-recognized epidemic among pregnant women. Clin. Obstet. Gynecol. 62, 168–184 (2019).

50. Hendrick, V. et al. Placental passage of antidepressant medications. Am. J. Psychiatry. 160, 993–996 (2003).

51. Rampono, J. et al. Placental transfer of SSRI and SNRI antidepressants and effects on the neonate. Pharmacopsychiatry. 42, 95–100 (2009).

52. Campbell, N. G., Koprich, J. B., Kanaan, N. M. & Lipton, J. W. MDMA administration to pregnant Sprague-Dawley rats results in its passage to the fetal compartment. Neurotoxicol. Teratol. 28, 459–465 (2006).

53. Pletscher, A. The 5-hydroxytryptamine system of blood platelets: physiology and pathophysiology. Int. J. Cardiol. 14, 177–188 (1987).

54. Brand, T. & Anderson, G. M. The measurement of platelet-poor plasma serotonin: A systematic review of prior reports and recommendations for improved analysis. Clin. Chem. 57, 1376–1386 (2011).

55. Anderson, G. M., Czarkowski, K., Ravski, N. & Epperson, C. N. Platelet serotonin in newborns and infants: Ontogeny, heritability, and effect of in utero exposure to selective serotonin reuptake inhibitors. Pediatr. Res. 56, 418–422 (2004).

56. Daws, L. C. Unfaithful neurotransmitter transporters: Focus on serotonin uptake and implications for antidepressant efficacy. Pharmacol. Ther. 121, 89–99 (2009).

57. Eddahibi, S. et al. Induction of serotonin transporter by hypoxia in pulmonary vascular smooth muscle cells. Relationship with the mitogenic action of serotonin. Circ. Res. 84, 329–336 (1999).

58. Zhang, H., Smith, G. N., Liu, X. & Holden, J. J. A. Association of MAOA, 5-HTT, and NET promoter polymorphisms with gene expression and protein activity in human placentas. Physiol. Genomics. 42, 85–92 (2010).

59. Li, Y. et al. GDM-associated insulin deficiency hinders the dissociation of SERT from ERp44 and down-regulates placental 5-HT uptake. Proc. Natl. Acad. Sci. U.S A. 111, E5697–E5705 (2014).

60. Blazevic, S. et al. Epigenetic adaptation of the placental serotonin transporter gene (SLC6A4) to gestational diabetes mellitus. PLoS One. 12, e0179934 (2017).

61. Schmon, B., Hartmann, M., Jones, C. J. & Desoye, G. Insulin and glucose do not affect the glycogen content in isolated and cultured trophoblast cells of human term placenta. J. Clin. Endocrinol. Metab. 73, 888–893 (1991).

62. Lang, I. et al. Human fetal placental endothelial cells have a mature arterial and a juvenile venous phenotype with adipogenic and osteogenic differentiation potential. Differentiation. 76, 1031–1043 (2008).

63. Livak, K. J. & Schmittgen, T. D. Analysis of relative gene expression data using real-time quantitative PCR and the 2-ΔΔCT method. Methods. 25, 402–408 (2001).

## Supplementary references

1. Naganuma, F. et al. Predominant role of plasma membrane monoamine transporters in monoamine transport in 1321N1, a human astrocytoma-derived cell line. J. Neurochem. 129, 591–601 (2014).

2. Van Lelyveld, N., Ter Linde, J., Schipper, M. E. I. & Samsom, M. Regional differences in expression of TPH-1, SERT, 5-HT3 and 5-HT4 receptors in the human stomach and duodenum. Neurogastroenterol. Motil. 19, 342–348 (2007).

3. Spandidos, A., Wang, X., Wang, H. & Seed, B. PrimerBank: A resource of human and mouse PCR primer pairs for gene expression detection and quantification. Nucleic Acids Res. 38, D792–D799 (2010).

4. Sun, Y. et al. Study of a possible role of the monoamine oxidase A (MAOA) gene in paranoid schizophrenia among a Chinese population. Am. J. Med. Genet. Part B Neuropsychiatr. Genet. 159 B, 104–111 (2012).

5. Métayé, T., Menet, E., Guilhot, J. & Kraimps, J. L. Expression and activity of G protein-coupled receptor kinases in differentiated thyroid carcinoma. J. Clin. Endocrinol. Metab. 87, 3279–3286 (2002).

